# Formation of Phage Lysis Patterns and Implications on Co-Propagation of Phages and Motile Host Bacteria

**DOI:** 10.1101/691568

**Authors:** Xiaochu Li, Floricel Gonzalez, Nathaniel Esteves, Birgit E. Scharf, Jing Chen

## Abstract

Coexistence of bacteriophages, or phages, and their host bacteria plays an important role in maintaining the microbial communities. In natural environments with limited nutrients, motile bacteria can actively migrate towards locations of richer resources. Although phages are not motile themselves, they can infect motile bacterial hosts and spread in space via the hosts. Therefore, in a migrating microbial community coexistence of bacteria and phages implies their co-propagation in space. Here, we combine an experimental approach and mathematical modeling to explore how phages and their motile host bacteria coexist and co-propagate. When lytic phages encountered motile host bacteria in our experimental set up, a sector-shaped lysis zone formed. Our mathematical model indicates that local nutrient depletion and the resulting inhibition of proliferation and motility of bacteria and phages are the key to formation of the observed lysis pattern. The model further reveals the straight radial boundaries in the lysis pattern as a tell-tale sign for coexistence and co-propagation of bacteria and phages. Emergence of such a pattern, albeit insensitive to extrinsic factors, requires a balance between intrinsic biological properties of phages and bacteria, which likely results from co-evolution of phages and bacteria.

**Author summary:** Coexistence of phages and their bacterial hosts is important for maintaining the microbial communities. In a migrating microbial community, coexistence between phages and host bacteria implies that they co-propagate in space. Here we report a novel phage lysis pattern that is indicative of this co-propagation. The corresponding mathematical model we developed highlights a crucial dependence of the lysis pattern and implied phage-bacteria co-propagation on intrinsic properties allowing proliferation and spreading of the microbes in space. Remarkably, extrinsic factors, such as overall nutrient level, do not influence phage-bacteria coexistence and co-propagation. Findings from this work have strong implications for dispersal of phages mediated by motile bacterial communities, which will provide scientific basis for the fast-growing applications of phages.

## Introduction

Viruses that specifically target bacteria, bacteriophages or phages, are critical components of the microbial world. They are found in almost every natural environment, including soil, waters, oceans, and bodies of macroorganisms (e.g., human guts) [1–3]. Furthermore, they are the most abundant organisms in the biosphere [2]. Through their interactions with bacteria, phages constantly regulate the ecology, evolution, and physiology of microbial communities [1,2]. Because of their antimicrobial activity, the application of phages in food processing, agriculture, and medicine has exploded in recent years [4–6]. Development of these applications benefits from fundamental knowledge about how phages interact with bacteria in a microbial community and how they are dispersed in their microenvironment.

As obligate parasites of bacteria, phages must coexist with their hosts at the population level [1]. This coexistence, however, appears rather inconceivable because phages have a huge proliferative advantage over bacteria. The generation cycles of phage and bacteria fall in comparable time frames, with both the phage latent period and bacterial division cycle on the order of an hour [7]. But in each generation cycle a bacterium produces two daughter cells, while one phage produces ~100 new phage particles. Thus, it would follow that phages would quickly outnumber and annihilate the host bacterial population [8,9]. However, phages and bacteria have coexisted in natural environments for eons. Recent theoretical and experimental studies demonstrated that the evolutionary arms race could maintain coexistence of phages with host bacteria [10–13]. Coevolution could drive a phenotypic and genotypic diversity in the ability of phages to attack the bacteria and the ability of bacteria to resist the attacks, thereby maintaining the balance between phages and host bacteria [10,14–16]. However, for a successful evolutionary arms race, phages and bacteria need to coexist at least over the time scale required for the emergence of beneficial mutations [8,9]. It is therefore critical to understand the population dynamics of phage-bacteria systems and conditions for their coexistence below the evolutionary time scale.

Previous studies on coexistence of phages and bacteria mostly focused on well-mixed, nearly homeostatic systems, such as cultures grown in chemostats [17–23]. Naturally occurring systems of phages and bacteria, however, often do not satisfy the conditions found under these defined laboratory settings. Firstly, natural systems typically do not offer a constant environment. Unlike chemostats, where steady levels of nutrients and waste are maintained, natural systems often experience sporadic deposition and replenishing of resources, and fluctuations in other conditions. Secondly, natural systems usually exhibit spatial heterogeneity to various degrees. The spatial inhomogeneity can significantly impact dynamical coexistence in the phage-bacteria systems [8,9,24–26].

A critical spatial process in the phage-bacteria system is the migration of bacteria and phages. Many motile bacteria can migrate towards nutrient-enriched areas via chemotaxis. Phages themselves are not motile, so their dispersal relies on either passive diffusion or transport by their hosts. However, diffusion is very inefficient for covering long distances. In addition, diffusion of phage particles is typically reduced by higher bacterial densities and increased viscosities due to bacterial exopolysaccharide production in biofilms [27–29]. Therefore, spatial dispersal of phages mostly relies on infection of and transportation by their motile host bacteria. It is poorly understood how phages and bacteria in a constantly migrating microbial community achieve coexistence, which implies their co-propagation in space.

In this work we explored the co-propagation of phages and motile bacteria using a simple experimental design, in which phages and bacteria were co-inoculated in a soft agar nutrient medium [30] (Fig 1a). The low agar concentration enabled motile bacteria to swim through the matrix, which, in combination with bacterial growth, resulted in the formation of visible “swim rings” [30]. Inoculation of bacteria and phages in separate locations allowed the experimental setup to mirror realistic scenarios, in which expanding bacterial populations encounter phages in a spatial domain. The described experiment generated a highly reproducible sector-shaped lysis pattern. This pattern cannot be explained by any previous mathematical models describing phage plaque formation [31–36], which inevitably produce circular patterns. Here we constructed a new mathematical model for the spatial dynamics of phages and bacteria, which reproduced the observed lysis pattern and revealed local nutrient depletion as the key to formation of the lysis pattern. Moreover, our model revealed that the sector-shaped lysis pattern with straight radial boundaries requires a balance between intrinsic biological properties of phages and bacteria, but does not depend on extrinsic factors. Such a pattern was further shown to be a tell-tale sign for extended spatial co-propagation of phages and bacteria, implying dependence of co-propagation on intrinsic balance between phages and bacteria. This is the first time that a sector-shaped lysis pattern has been reported in phage-bacteria systems. Our study of this phenomenon via an integrated modeling and experimental approach provides critical insights into naturally occurring dispersal and cohabitation of phages infecting motile bacteria.

**Figure 1.**
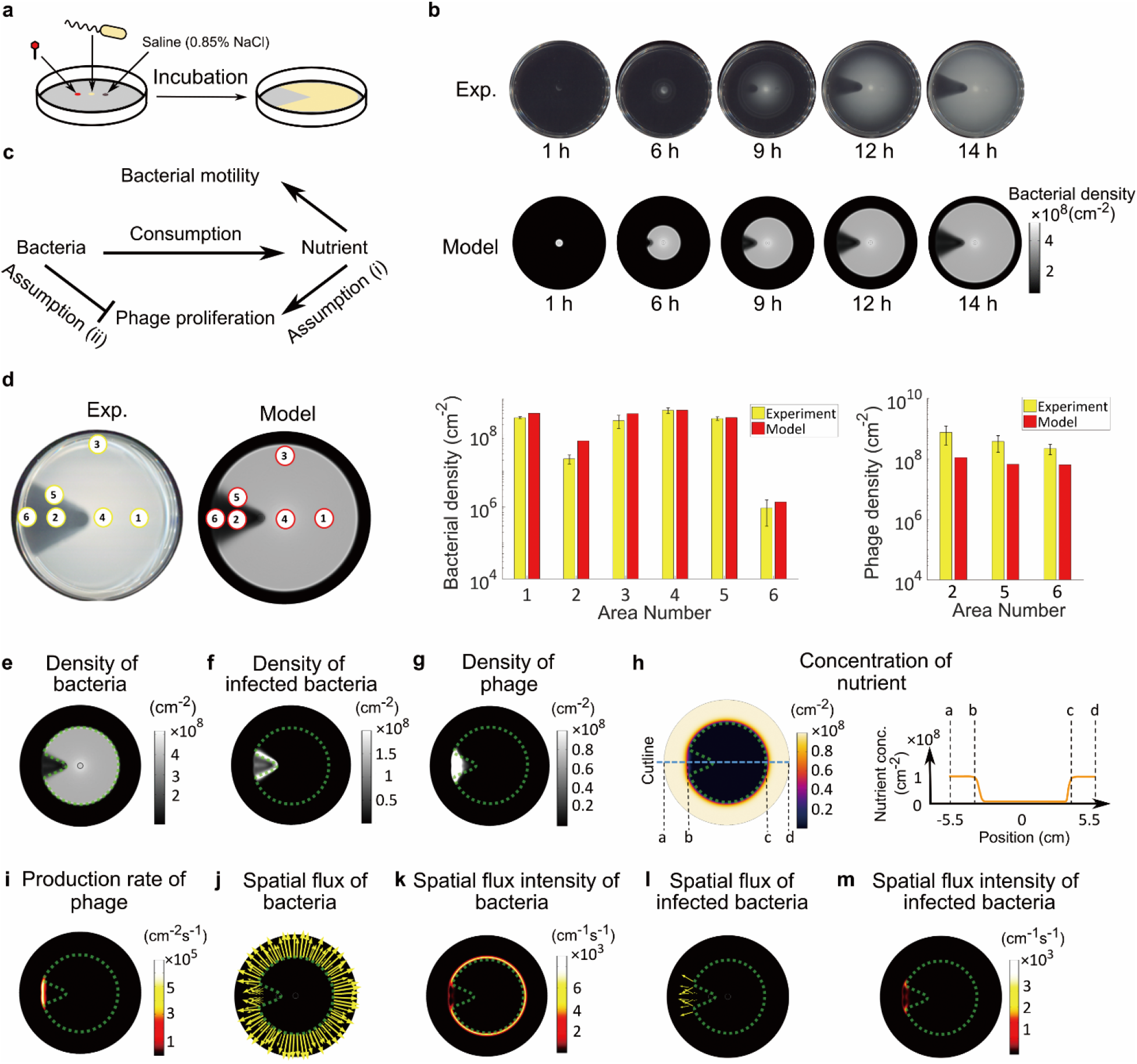
Sector-shaped lysis patterns emerge due to nutrient depletion. (a) Schematic of experimental procedure. Bacteria, phages, and a saline control were spotted on swim plates and incubated for 14 hours, which resulted in the development of the sector-shaped lysis pattern. (b) Experimental and model results of the lysis pattern over time. (c) Interactions between key processes in the model. Pointed arrows: positive influences. Blunt arrows: negative influences. (d) Quantitative comparison of bacterial and phage densities between experiment (yellow bars) and model (red bars). In the experiment, areas labeled by yellow numbers were sampled for phage and bacteria quantifications (see Methods). Corresponding areas in the model are labeled by red numbers. (e) Simulated density of total bacteria. The dashed green outline of the bacteria-dense area is superimposed on (f-m) for reference. (f) Simulated density of infected bacteria. (g) Simulated density of phages. (h) Simulated nutrient concentration (per unit area). Right: Orange curve shows the nutrient concentration profile along the axis of symmetry of the lysis pattern (blue dashed line in left panel). (i) Simulated phage production rate. Active phage production only happens at the outer edge of the lysis area. (j) Simulated spatial flux of total bacteria. (k) Intensity of bacterial spatial flux (~ length of arrow in (j)). Bacteria are motile only at the outer edge of the swim ring. (l) Simulated spatial flux of infected bacteria. (m) Intensity of spatial flux of infected bacteria (~ length of arrow in (l)). Infected bacteria are only motile at the outer edge of the lysis area. The spatial flux shown in (j-m) represents the sum of diffusion flux and chemotaxis flux. Length of the arrow is proportional to magnitude of the spatial flux. (e-m) present snapshots of model simulation at 10 h, an intermediate time at which the colony expansion and pattern formation progress steadily.

## Results

### Nutrient depletion is critical for formation of the lysis pattern

We designed a series of quantitative experiments based on our previously described phage drop assay [41], which allowed a spatially propagating bacterial population to encounter phages. *Salmonella enterica* serovar typhimurium 14028s and χ phage were inoculated 1 cm apart (Fig 1a) on 0.3% agar plate containing bacterial growth medium [30,42]. As the bacterial population grew, nutrients were consumed. Due to the low agar concentration, bacteria swam through the matrix and followed the self-generated nutrient gradient via chemotaxis, causing spreading of the bacterial population and the appearance of a swim ring. As the bacterial swim ring expanded, it reached the phage inoculation point. The phages then infected the bacteria and generated a lysis area with low bacterial density in the swim ring (Fig 1b). This experiment gave rise to an intriguing sector-shaped lysis pattern (Fig 1b). Most strikingly, as the bacterial swim ring expanded, the radial boundaries of the lysis area stayed unchanged behind the expanding front, resulting in a frozen or immobilized lysis pattern (Fig 1b, S1 Movie). Once the spreading of the swim ring stopped at the plate wall, the lysis pattern persisted for at least 48 hours (data not shown).

To understand the formation of this lysis pattern, we constructed a mean-field partial differential equation (PDE) model for the phage-bacteria system (Eqs.(1) ~ (4)). Like the previous phage plaque models [31–36], our model depicted the basic processes underlying the proliferation and propagation of phages and bacteria. Namely, the bacteria consume nutrients, divide, and move up the nutrient gradient via chemotaxis-directed swimming motility. Once infected by phages, the bacterium is lysed after a latent period, and release new phage progeny. Note that the run-and-tumble mechanism of bacterial chemotaxis results in a biased random walk of cells up the nutrient gradient. The random walk was expressed in the model as the cell diffusion terms and the bias as the cell drift terms (Eqs.(1) and (2)). In addition, we incorporated the following new assumptions about phage-bacteria interactions in the model, which were critical elements for formation of the sector-shaped lysis pattern (S1 Fig).

i. Nutrient deficiency inhibits phage replication (Fig 1c). Because phage replication in the host bacteria requires energy, it is likely reduced at low nutrient levels.
ii. High bacterial density inhibits phage production (Fig 1c). Such an effect has previously been implied in *Escherichia coli* and phage λ [43]: *E. coli* cells reduce the number of phage receptors in response to externally applied quorum sensing signals, resulting in decrease of phage adsorption rate and ultimately overall phage production.

Our model was able to reproduce the lysis pattern observed in the phage drop assay (Fig 1b, S1 Movie) and quantitatively match the bacterial and phage density profiles throughout different areas of the agar plate (Fig 1d). Both, experimental and modeling results, displayed the highest bacterial density at the inoculation point (area 4), followed by areas outside the lysis sector (areas 1 and 3), the radial boundary of the lysis pattern (area 5), and in the middle of the lysis pattern (area 2). The lowest bacterial density was at the outer edge of the lysis sector (area 6). The predicted phage densities also matched the experimental values, with the highest number of phages localized in the middle of the lysis sector and the lowest number present at the edge of the swim ring near the wall of the plate (Fig 1d). It should be noted that the lysis area was not entirely void of bacteria. In both experimental and modeling results, a low density of bacteria remained within the lysis area. In the model, nearly all bacteria in this area were infected bacteria (Fig 1e & f). In reality, this subpopulation could also include phage-resistant bacteria, which was not encompassed in our current model.

The model results revealed local nutrient depletion as the key reason for the lysis pattern to immobilize behind the expanding front of the bacterial swim ring. As the swim ring expanded, nutrients were depleted within the ring (Fig 1h). Nutrient depletion inhibited both phage production (Fig 1i) and bacterial motility (Fig 1j-m). Note that phages relied on the infection of motile bacteria to propagate in space. The passive diffusion of phage particles (*D_P_* ~ 1 *μm*^2^*h*^−1^) was negligible compared to the effective diffusion of bacteria resulting from run-and-tumble (*D_B_* ~ 10^5^*μm*^2^*h*^−1^) (Table 1). Therefore, inhibition of bacterial motility, especially motility of the infected bacteria (Fig 1l & m), also hindered spatial propagation of phages. Together, the inhibition of phage production and propagation due to local nutrient depletion and reduction of bacterial motility resulted in immobilization of the lysis pattern at the interior of the bacterial swim ring. The lysis pattern only actively grew at the expanding front of the swim ring, where nutrient supply from the unoccupied periphery supported active phage production and propagation (Fig 1i-m).

**Table 1:**
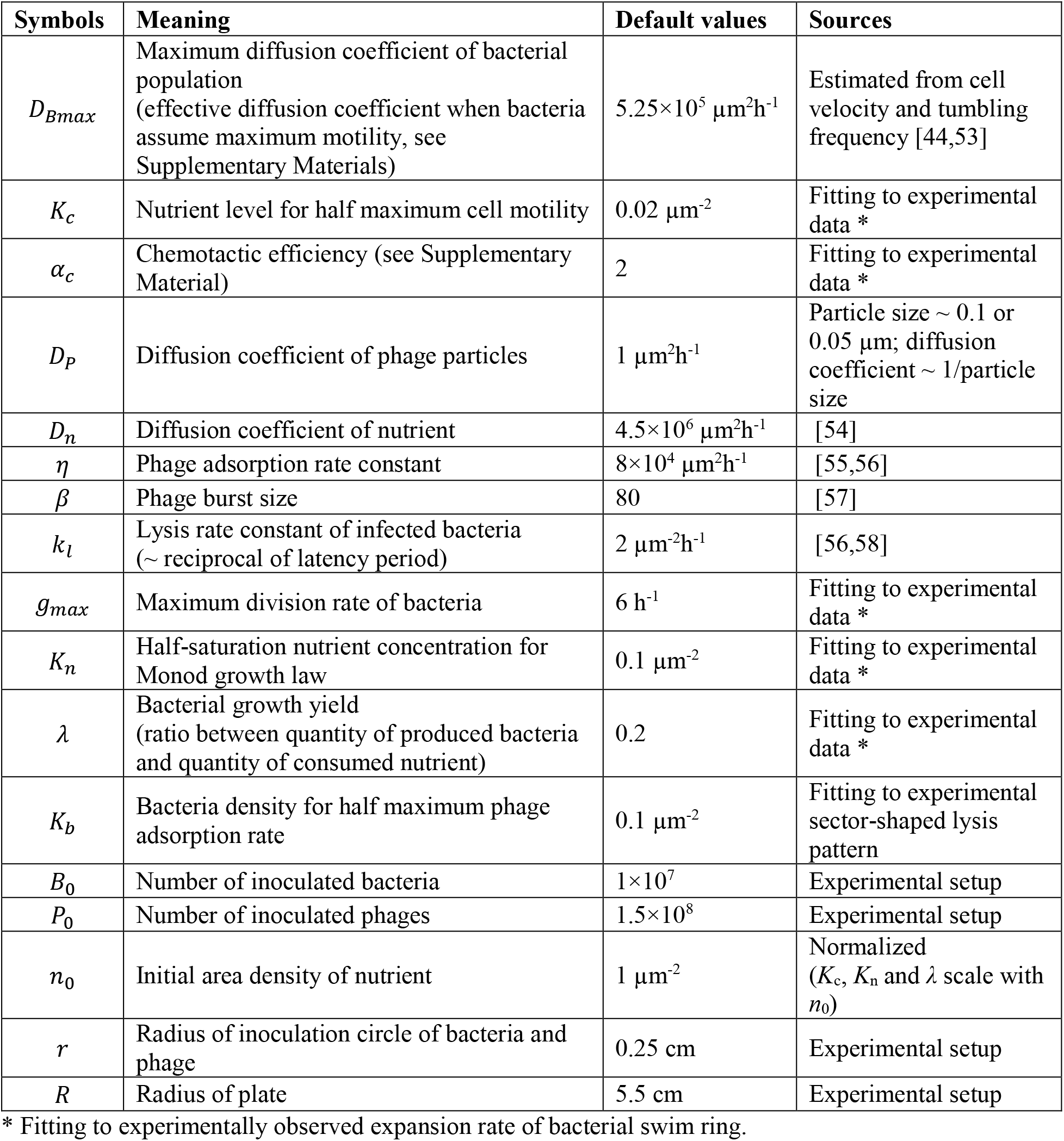
Parameters of mathematical model.

### The lysis pattern reflects radial projection of phage initiation zone

Interestingly, the experiment presented a decrease in the angle of the lysis sector when phages were inoculated further away from the bacterial inoculation point and vice versa (Fig 2a). This observation was faithfully reproduced by our model (Fig 2a). The lysis patterns in these cases approximately reflected the radial projection from the bacterial inoculation point over an approximately 0.7 cm circle centered at the phage inoculation point (Fig 2a, cartoon). In the model, we found that this projected area roughly corresponded to the area occupied by phages when nutrients initially got depleted at the phage inoculation point (Fig 2a, 3^rd^ and 4^th^ rows, nutrient depleted to 5 % of initial level). We hereby termed this area the “phage initiation zone”. The phage initiation zone marked the initialization of the steady expansion of the lysis pattern. After the phage initiation zone was established, the phage and bacterial densities at the expanding front remained at a steady level throughout the rest of the pattern formation (S2 Fig).

**Figure 2.**
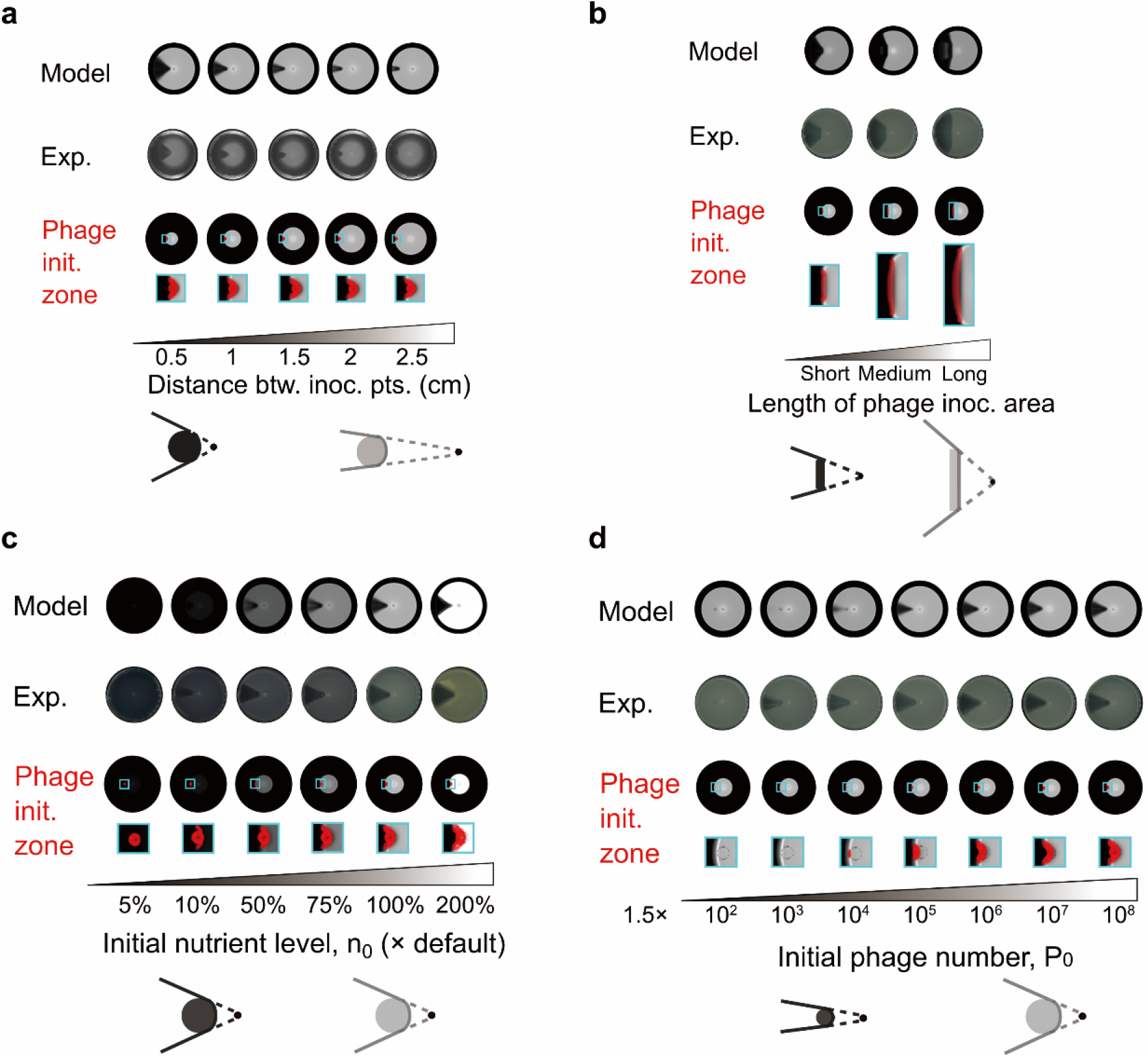
The lysis pattern reflects the radial projection of the phage initiation zone and is insensitive to extrinsic factors. Model and experimental results with (a) phages inoculated at different distances from the bacterial inoculation point, (b) phages inoculated with rods of different lengths, (c) different initial nutrient levels, and (d) different initial phage number. In (a-d), the phage initiation zones represent areas occupied by phages when the nutrient level at the center of the phage inoculation area drops below a certain threshold (set at 5 % of the initial nutrient level in model simulations). Cartoons summarize factors leading to different lysis patterns. Black dots: bacterial inoculation point. Solid dark grey and light grey areas: illustrative phage initiation zones. Solid lines: radial boundaries of lysis patterns. Dashed lines: extension from bacterial inoculation points to boundaries of the lysis areas.

Our model further predicted how the phage initiation zone and the projected lysis pattern rely on additional factors. The predictions were all confirmed by experiments (Fig 2). Firstly, the phage initiation zone encompassed the original phage inoculation area (Fig 2a & b). Particularly, when phages were inoculated with rods that were significantly longer than the size of the phage initiation zone during point inoculation, the phage initiation zone became dominated by the rod size, and the lysis pattern roughly reflected the radial projection of the phage inoculation area (Fig 2b). Secondly, the size of the phage initiation zone remained roughly the same despite changes in total nutrient concentrations, resulting in similar angles of the lysis sector (Fig 2c). Thirdly, the phage initiation zone enlarged as initial phage particle number increased, resulting in a larger angle in the lysis sector (Fig 2d). Lower phage inoculation density led to decreased phage production and propagation by the time nutrient got depleted locally by bacteria. When initial phage numbers were too low, phages failed to establish the initiation zone, and subsequently, the lysis sector (Fig 2d, *P*_0_= 1.5 × 10^2^). The model also predicted that the initial bacteria number does not affect the lysis pattern (S3 Fig). Remarkably, the lysis pattern maintained straight radial boundaries (Fig 2) despite changes in the extrinsic factors tested above, i.e., distance between inoculation point, size of inoculation area, overall nutrient level, and initial phage/bacteria number.

### Competition between phages and bacteria determines shape of lysis pattern

Although the straight radial boundaries of the lysis pattern were maintained under various external conditions like nutrient level and initial inoculation, our model predicted a significant change in the lysis pattern when intrinsic biological parameters were altered. In the simulation results, promoting the proliferative efficiency of phages (e.g., by increasing phage adsorption rate, phage burst size and lysis rate of infected bacteria) caused the lysis pattern to flare out, and decreasing phage proliferation caused the lysis pattern to close up (Fig 3, horizontal axes). Meanwhile, promoting bacterial proliferation caused the lysis pattern to curve inward and close up, and vice versa (Fig 3, vertical axes). In detail, the model assumed dependence of both bacterial and phage proliferation on nutrient (both are increasing functions of local nutrient concentration, see Materials and Methods). As previously shown, proliferation of bacteria caused nutrient to be depleted inside the bacterial swim ring (Fig 1h). Therefore, the bacterial proliferation rate at the expanding front determined how fast nutrients were depleted locally.

**Figure 3.**
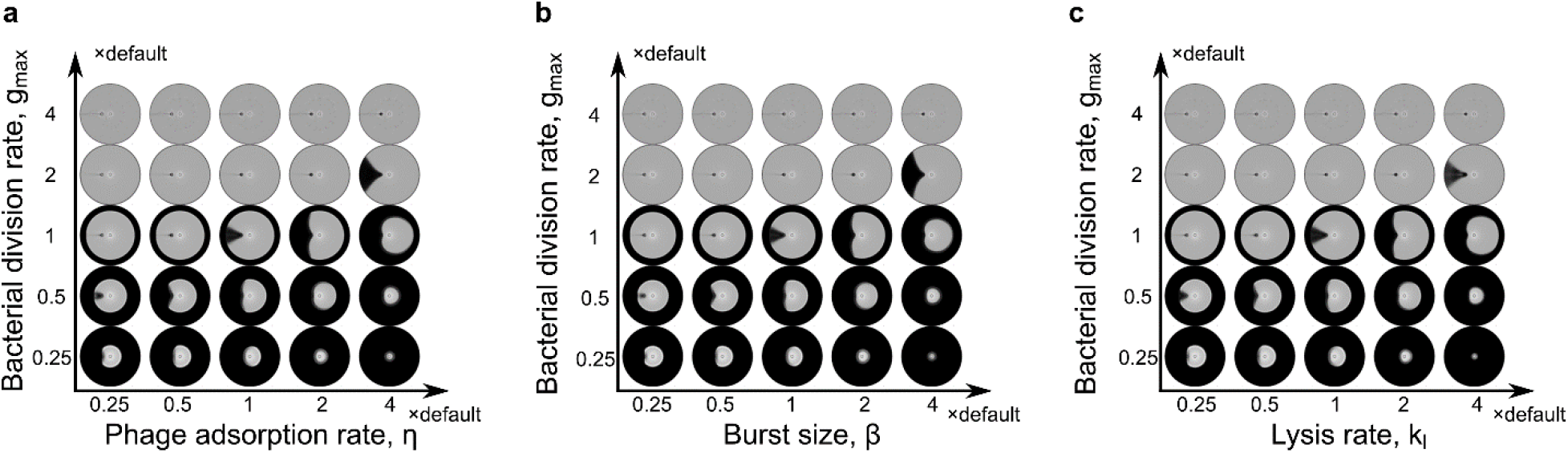
Competition between bacterial and phage proliferation determines the shape of lysis patterns. Simulated lysis patterns with various bacterial growth rate constants versus (a) phage adsorption rate constants, (b) phage burst sizes, and (c) phage-induced lysis rate constants.

This time further determined angular spreading of phages along the expanding front of the swim ring, because phage proliferation only thrived before nutrient was depleted locally. Therefore, either stronger phage proliferation or a weaker bacterial proliferation (causing slower nutrient consumption) allowed phages to spread in an accelerated fashion as the swim ring expanded, resulting in a flared-out lysis pattern. Vice versa, a weaker phage proliferation relative to bacterial proliferation resulted in a lysis pattern with edges closing inwards. The experimentally observed sector-shaped lysis pattern maintained straight radial boundaries despite changes in extrinsic variables, indicating that this observed pattern stemmed from a balance between proliferation of the bacterial and phage strains tested.

### Bacterial motility and chemotaxis affects the lysis pattern

We next used the model to investigate how bacterial motility and chemotaxis influenced the shape of the lysis pattern. Bacterial motility was reflected by the bacterial diffusion coefficient in the model. A larger diffusion coefficient corresponds to higher cell speed [44]. Expectedly, a larger diffusion coefficient caused faster expansion of the bacterial swim ring in the model (Fig 4a). The chemotactic efficiency, on the other hand, characterized the bias of diffusion. Chemotaxis promoted the directed motility of bacteria in the radial direction due to the nutrient gradient formed by bacterial nutrient consumption (Fig 1h). Consistently, the model predicted that higher chemotactic efficiency expedites expansion of the bacterial swim ring (Fig 4a), because the moving cells at the expanding front can follow the nutrient gradient more efficiently.

**Figure 4.**
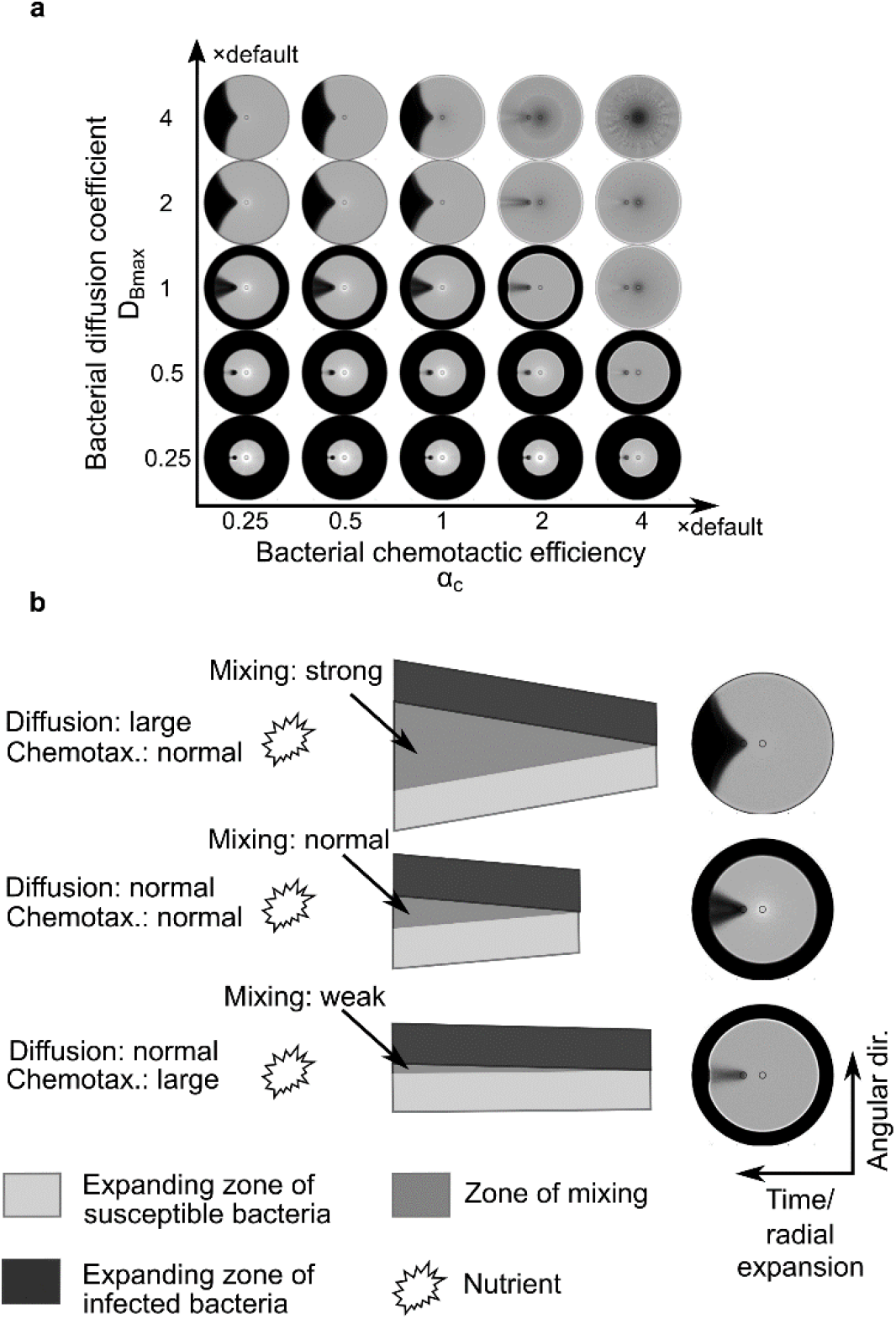
Effects of bacterial motility and chemotaxis on the shape of lysis patterns. (a) Simulated lysis patterns with various bacterial diffusion coefficients and chemotactic efficiencies. The diffusion coefficient in the model reflects the efficiency of bacterial motility. (b) Illustration of how bacterial diffusion and chemotaxis affect the evolution of lysis patterns. The cartoon illustrates a hypothetical history of expansion and mixing of two bacterial patches that would occur along the expanding front of the swim ring. Dark and light grey patch starts with infected and susceptible bacteria, respectively. Because nutrient is depleted behind the expanding front, bacterial expansion along the radial axis only occurs in the outward direction. Large bacterial diffusion promotes expansion equally in all directions, which enhances mixing between the infected and susceptible bacteria and leads to more effective phage propagation and flare-out of the lysis pattern (top row). In contrast, large chemotactic efficiency promotes expansion only against the nutrient gradient (radial direction), which effectively parallelizes bacterial motion, reduces their mixing in the angular direction, and causes the lysis pattern to close up (bottom row).

The effects of bacterial motility and chemotactic efficiency on the lysis pattern, however, were predicted to be exactly opposite to each other. According to the simulation results, increasing the bacterial diffusion coefficient or decreasing the chemotactic efficiency caused the pattern to flare out (Fig 4a). Vice versa, decreasing the bacterial diffusion coefficient or increasing chemotactic efficiency caused the pattern to close up (Fig 4a). The model generated this result because increasing bacterial diffusion coefficient, i.e., increasing bacterial motility, promoted mixing of infected and susceptible bacteria (Fig 4b, top row). Such mixing was critical for spatial propagation of phages and angular expansion of the lysis pattern, because phages could not move on their own and relied on infected bacteria to spread in space. In contrast, enhancing chemotaxis inhibited such mixing, because it effectively promoted parallel motion of the bacteria along the radial direction towards the high-nutrient area outside the swim ring (Fig 4b, bottom row). Taken together, bacterial motility and chemotaxis needed to be in balance to generate a lysis pattern with straight radial boundaries. Collectively, these findings and those from the model in the previous section indicated that bacterial motility and chemotaxis are required to be in balance with bacterial and phage proliferation rate to generate straight radial boundaries in the lysis pattern (S4 Fig).

To test the model predictions, we performed the phage drop assay with strains of *S*. typhimurium 14028s containing deletions in two chemoreceptor encoding genes, *tar* and *tsr*. Strains with deletions in *tar* or *tsr* did not significantly change the lysis pattern, whereas the strain containing a deletion of both genes produced a moderate flare-out of the lysis pattern (S5 Fig). This result was qualitatively consistent with the model prediction that weaker chemotactic efficiency caused a pattern with a wider angle (Fig 4a).

### Straight radial boundary of lysis pattern is a tell-tale sign for extended co-propagation

A closer look at the model results revealed that the straight lysis pattern boundaries in the lysis pattern implied co-propagation of bacteria and phages over extended periods. Unlike the sector-shaped pattern with straight radial boundaries, a flared-out or closed-up lysis pattern indicated that one species would outcompete the other during the co-propagation (Fig 5a). For example, the result at the upper right corner of Fig 5a shows a case where phages encircled bacteria and blocked their further propagation in space. Vice versa, the result at the lower left corner of Fig 5a shows the opposite case where phage propagation was blocked by bacteria. For the less extreme flared-out or closed-up lysis patterns (e.g. upper left and lower right corners of Fig 5a), one species would eventually encircle and block the other, if the simulation had been run on a larger spatial domain that allowed further spatial expansion.

**Figure 5.**
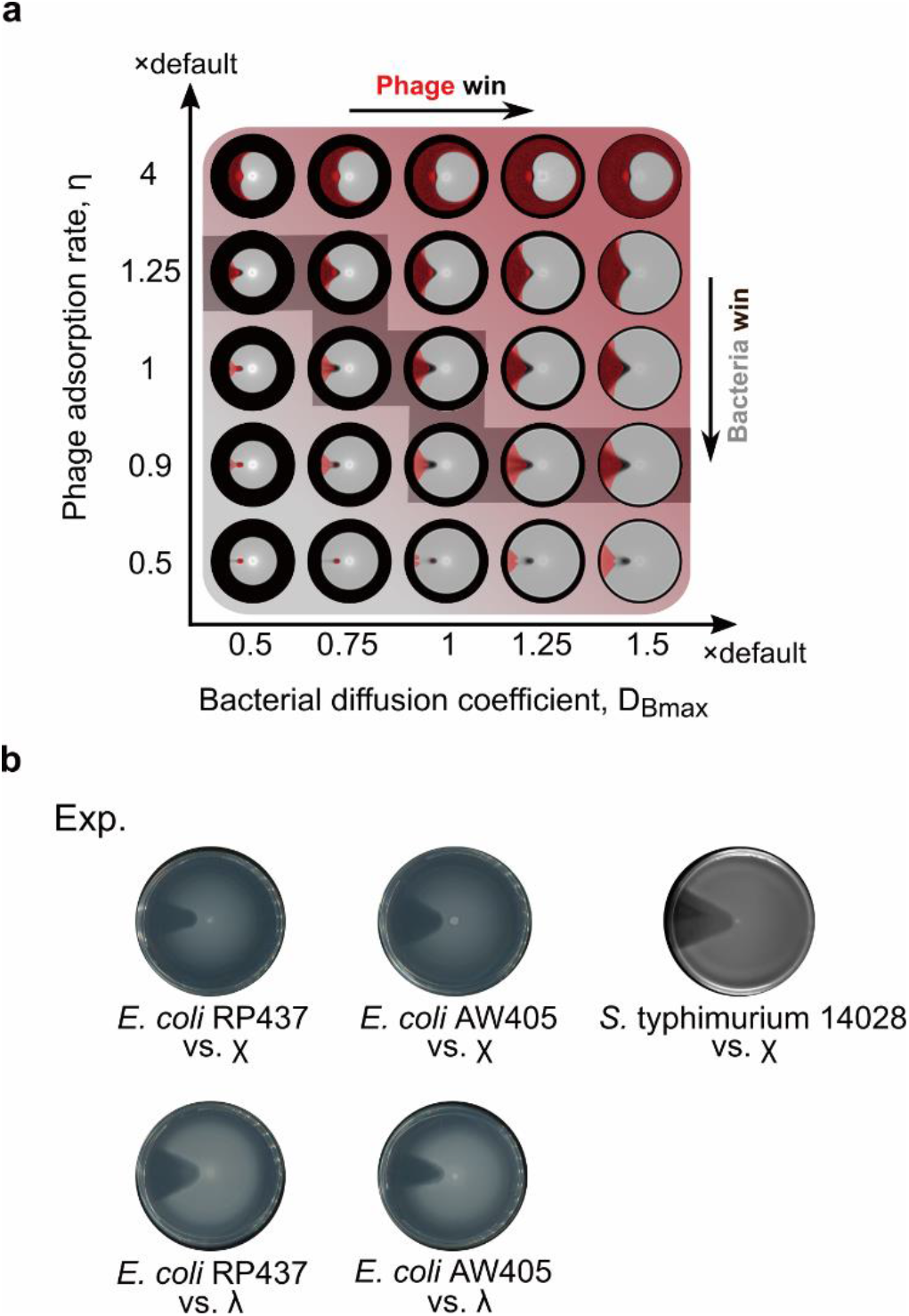
Co-propagation between phage and bacteria is reflected by the straight radial boundary of lysis pattern. (a) Lysis patterns and corresponding distribution of phages with various bacterial diffusion coefficients and phage adsorption rate constants. Red shade: density of phages. Grey shade: density of bacteria. Grey shadow staircase: potential trajectory of evolutionary arms race. (b) Experimental results from pairs of different bacterial species and their cognate phages.

Remarkably, similar sector-shaped lysis patterns with straight radial boundaries were observed in different bacteria-phage pairs (Fig 5b). First, we performed the phage drop assay using *E. coli* and phage χ, and found a similar sector-shaped lysis pattern (Fig 5b, first row). Since χ is a bacterial flagella-dependent phage [45–47], we further examined whether infection of motile *E. coli* by a non-flagellotropic phage, λ, would generate a similar lysis pattern. Phage λ was chosen because of an overlapping bacterial host range with χ. Interestingly, similar sector-shaped lysis patterns were observed independent of the utilized phage type (Fig 5b). Combined with the model findings above and earlier results on sensitivity of the pattern to intrinsic parameters, these highly similar sector-shaped lysis patterns indicated that both *Salmonella* and *E. coli* achieved intrinsic biological balance with both phages, λ and χ.

## Discussion

In this work, we combined experiments and modeling of lysis pattern formation to investigate the coexistence and co-propagation of phages and bacteria in space. Our experimental setting has strong implications for realistic scenarios, where an expanding bacterial population encounters phages and mediates their dispersal. We observed the formation of an asymmetric, sector-shaped lysis pattern, which cannot be explained by previous models for lysis pattern formation in phage-bacteria systems. Our new mathematical model successfully reproduced the experimental observation and revealed the importance of nutrient depletion in maintaining the geometric asymmetry initialized in the system. Specifically, local nutrient depletion inhibited phage production and propagation behind the expanding front of the bacterial swim ring, thus immobilizing the lysis pattern. Without the immobilization effect, the lysis pattern would lose asymmetry and eventually reduce to a circle, as predicted by the previous models for phage plaque formation [31–36].

Most importantly, straight radial boundaries of the lysis pattern presented a tell-tale sign that a phage-bacteria system is capable of co-propagation over extended period (Fig 5a). Therefore, the shape of the lysis pattern can serve as a reporter of co-propagation. The model further demonstrated that a straightly expanding pattern requires balance between bacterial and phage proliferation efficiency, and bacterial motility and chemotaxis (Figs 3, 4 and S4 Fig). The balance of biological properties keeps angular expansion of the lysis pattern in pace with its radial expansion and creates straight radial boundaries in the evolving lysis pattern. In contrast, the straight boundary is insensitive to extrinsic factors, such as nutrient levels and initial phage/bacterial numbers (Fig 2, S3 and S6 Figs). Together, these findings suggest that balance of intrinsic factors supports robust co-propagation of phages and bacteria regardless of variations in the environment or initial conditions. Interestingly, we experimentally discovered similar sectorial lysis patterns with straight radial boundaries in two different enteric bacterial species paired with two different phages (Fig 5b). These lysis patterns implied intrinsic biological balance between the phages and their bacterial hosts. This phenomenon suggests that natural pairs of bacteria and phages could have shaped their biological properties to allow robust spatial coexistence and co-propagation, likely as a result of coevolution. In the future, we will perform the phage drop assay on other phage-bacteria pairs to examine whether this conclusion may be generalized universally.

The model predictions on extrinsic factors were verified by our experiments (Fig 2). It was much more complicated, though, to vary the intrinsic biological properties in a controlled fashion. We are relegating these experimental testing to future work, which will also provide feedback for model refinement. For example, the predicted lysis pattern only changed significantly when chemotactic efficiency was increased from the default value (Fig 4a), whereas a flared-out lysis pattern occurred experimentally in a strain lacking two major chemoreceptors (S5 Fig). This quantitative discrepancy indicates that the chemotaxis term and/or parameters in the model should be modified in the future.

Our study specifically underscores the importance of co-propagation of phages and bacteria, i.e., their coexistence in the context of a migrating microbial community. We found that co-propagation requires not only a balance between the proliferative efficiency of phages and bacteria, but also between their ability to spread in space (autonomous spreading of bacteria vs. bacteria-mediated spreading of phages). The requirement of balanced proliferative efficiency is known to create the selective pressure that drives the evolutionary arms race between phages and bacteria in their ability to attack and resist attack [10,12,14–16]. Similarly, the requirement of balanced ability to spread in space could also create a selective pressure to drive an arms race in evolving stronger ability to spread in space (Fig 5a, dark shaded staircase). For example, the emergence of flagellotropic phages, i.e., phages specifically targeting actively rotating bacterial flagella [48], could reflect an evolved strategy for phages to improve their ability to propagate in space. Interestingly, our model demonstrates a significant impact of chemotactic efficiency on co-propagation of phages and bacteria (Fig 4 and S6d Fig). Therefore, bacteria could theoretically evolve higher chemotactic efficiency as a counterattack on phage infection. In the future we will examine whether the arms race between phages and bacteria indeed affect the diversity in genes regulating bacterial chemotaxis.

Our current work exploited the simplest possible experimental and model setup to understand how phages and bacteria coexist and co-propagate in space, using lytic phages and uniform initial nutrient concentration. In the future, we will modify our experimental and model parameters to investigate additional factors, such as lysogeny and non-uniform nutrient distribution, on the spatial dynamics of phages and bacteria. We will also incorporate coevolution between phages and bacteria into our model and experimental set up, to investigate the long-term co-propagation under the effect of evolution. Findings from this work have strong implications for dispersal of phages in microbial communities and lay the groundwork for future applications, such as phage therapy. Ultimately, we hope to create a model that will aid successful selection and engineering of phages for targeted applications by providing information on phage dispersal and interaction with host bacteria in the corresponding environment.

## Materials and Methods

### Bacterial strains and phages

The strains of bacteria and phages used are listed in Table 2.

**Table 2:** Biological materials used in this study.

| Species/strains/<br>plasmids | Relevant characteristics | Sources | References |
| --- | --- | --- | --- |
| *Salmonella enterica* serovar typhimurium |  |  |  |
| 14028s | Wild type | Gift from Rasika M. Harshey |  |
| 14028s Tar<sup>-</sup> | *tar<sup>-</sup>* | This work |  |
| 14028s Tsr<sup>-</sup> | *tsr<sup>-</sup>* | This work |  |
| RH2312/SM20 | *tar<sup>-</sup>tsr<sup>-</sup>* | Gift from Rasika M. Harshey |  |
| TH2788 | *fliY5221::Tn10dTc* (-86 from ATG of *fliY*) | Gift from Kelly T. Hughes |  |
| *Agrobacterium* sp. H13-3 |  |  |  |
| RU12/001 | Sm<sup>r</sup>; spontaneous streptomycin resistant wild-type strain |  | [49] |
| *Escherichia coli* |  |  |  |
| RP437 | Wild type | Gift from Howard C. Berg | [50] |
| AW405 | Wild type | Gift from Howard C. Berg | [51] |
| Phages |  |  |  |
| λ phage | *vir* | Gift from Rüdiger Schmitt | [52] |
| χ phage |  | Gift from Kelly T. Hughes |  |
| Plasmid |  |  |  |
| pKD46 | *bla* P<sub>BAD</sub> *gam* *beto* *exo* pSC101 *oriTS* | Gift from Howard C. Berg | [37] |

### Media and growth conditions

*Salmonella enterica* serovar Typhimurium 14028s was grown in MSB at 37 °C. MSB is a modified LB medium (1 % tryptone, 0.5 % yeast extract, and 0.5 %NaCl) supplemented with 2 mM MgSO_4_ and 2 mM CaCl_2_. *Escherichia coli* strains were grown at 30 °C in T-broth containing 1 %tryptone and 0.5 %NaCl at 30 °C.

### Construction of mutant strains

The protocol for lambda-Red genetic engineering [37,38] was followed to make *S*.typhimurium mutant lacking *tar*.

### Phage drop assay

Swim plates containing MSB medium (for *S*. Typhimurium) or T-broth (for *E. coli)* and 0.3%bacto agar were inoculated with 2.5 μl of a stationary phase bacterial culture in the center of the plate along with 2.5 μl of phage suspension (MOI = 25.4) at a 1 cm distance from the inoculation point. A 2.5 μl spot of 0.85% saline was placed at the same distance from the bacterial inoculation point, opposite from the phage suspension, as a control. pPlates were incubated at 37 °C (S. Typhimurium) or 30 °C (*E. coli*) for 14 hours. All plates were imaged using the Epson Perfection V370 scanner. Phage drop assays with slight modifications were conducted to test different variables. For the rod shaped inoculations, sealed glass capillaries of different lengths were immersed in phage suspension and pressed against the soft agar at a distance of 0.5 cm from the bacterial inoculation point. To test the effect of varying inoculation distances, phage suspensions were inoculated at 0.5, 1.0, 1.5, 2.0, and 2.5 cm from the bacterial inoculation point. For altered nutrient concentration experiments, the initial concentrations of tryptone and yeast extract were adjusted to be 0.05, 0.1, 0.5, 0.75, or 2.0 times of the regular nutrient concentration, which is referred to as a concentration of 1. To evaluate the effect of phage number, the initial phage stock was serially diluted ten-fold and then spotted on the plate. In experiments conducted with λ phage, the swim plates were supplemented with 10 mM MgSO_4_ and 0.2 % maltose.

### Phage titer

Serial dilutions of the phage stock were made and 100 μl of each dilution was added to host bacterial cells with an OD_600_ of 1.0. Bacteria-phage mixtures were incubated for 10 min at room temperature. Each mix received 4 ml of pre-heated 0.5 % soft agar and was then overlaid on LB plates. Plates were incubated at 37 °C for 4-6 h. The titer of the phage stock was determined by counting the plaques on the plate that yielded between 30 to 300 plaque forming units and multiplying the number by the dilution factor.

### χ phage preparation

Dilutions of phage suspensions mixed with bacteria were plated to achieve confluent lysis as described in the phage titer protocol using 0.35 % agar for the overlay. Following formation of plaques, 5 ml of TM buffer (20 mM Tris/HCl [pH=7.5], 10 mM MgSO_4_) was added to each plate and incubated on a shaking platform at 4 °C for a minimum of 6 h. The soft agar/buffer mixture was collected, pooled, and bacteria were lysed by adding chloroform to at final concentration of 0.02 %. Samples were mixed vigorously for 1 min, transferred to glass tubes, and centrifuged at 10,000 x g for 15 min at room temperature. The supernatant was passed through a 0.45 μm filter and NaCl was added to a final concentration of 4 %. The protocol of phage preparation was followed as described in [39]. The final phage stock was stored in TM buffer at 4 °C.

### Bacteria and phage quantifications

Phage drop assays were conducted as described above. At the 14-hour end point, different areas of the plate were sampled by taking agar plugs using a 10 ml syringe barrel with plunger. Each agar plug was placed in 1 ml of 0.85 % saline and incubated at room temperature for 10 min with shaking to allow even mixture of the agar. Serial dilutions of each sample were plated on LB agar plates and incubated at 37 °C overnight. For phage quantifications, 100 μl of chloroform was added to each sample. The number of phage particles present in each sample was quantified as described in the phage titer protocol. Densities reported correspond to plaque forming units (for phage) or colony forming units (for bacteria). To compare with model results, the volume densities were converted to area densities, based on 0.5 cm thickness in the agar, i.e., area density (cm^−2^) = volume density (CFU/cm^3^ or PFU/cm^3^) × 0.5 cm.

### Model setup

We constructed a mean-field diffusion-drift-reaction model for the bacteria-phage system. Our model includes four variables: density of susceptible bacteria *B*(*x, t*), density of infected bacteria *L*(*x, t*), density of phages *P*(*x, t*), and nutrient concentration *n*(*x, t*). The state variables and parameters are summarized in Table 1. The equations governing the spatiotemporal dynamics of bacteria, phages and nutrient read as Eqs. (1) ~ (4).

Susceptible bacteria:

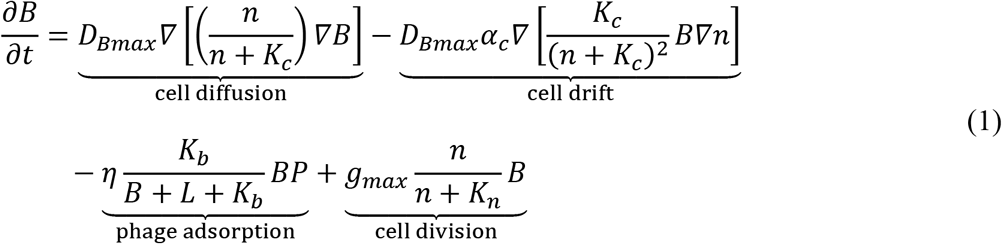

Infected bacteria:

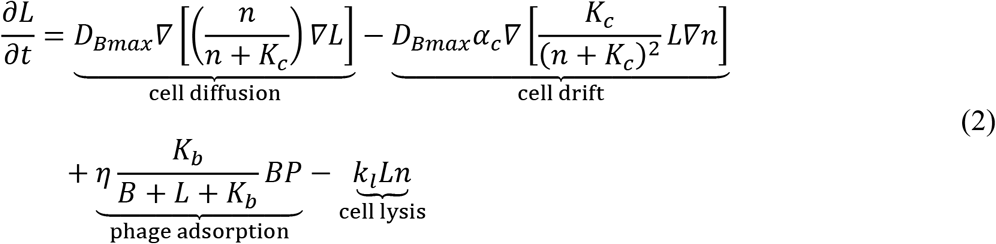

Phages:

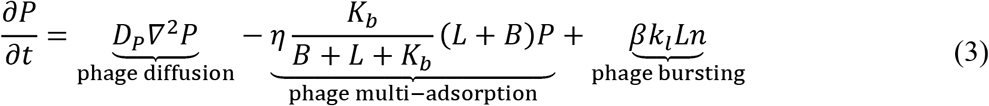

Nutrient:

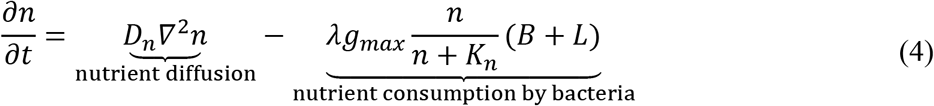

Eqs. (1) ~ (4) incorporate the following model assumptions.

1. The division rate of susceptible bacteria follows the Monod rate law [40].
2. Division of the infected bacteria is neglected because they are likely lysed before dividing. But they consume nutrients at the same rate as susceptible bacteria (changing this rate does not affect the qualitative behavior of the model).
3. The phage adsorption rate decreases with increasing bacterial density. This is how New Assumption (ii) in Results is implemented.
4. Multi-adsorption is considered, i.e., phages can be adsorbed onto bacteria that are already infected.
5. Because phage assembly requires energy, we assume that the lysis period elongates as nutrient level decreases. This is how New Assumption (i) in Results is implemented.
6. Because bacterial motility requires energy, it depends on nutrient level. This dependence is reflected in both the diffusion and chemotaxis terms. Derivation of the diffusion and chemotaxis terms is given in the Supplementary Materials and S7 Fig. Both susceptible and infected bacteria follow these spatial dynamics.

## Acknowledgements

We thank Kelly T. Hughes for the gift of χ phage, Rüdiger Schmitt for λ phage, Howard C. Berg for providing us with pKD46 and various *E. coli* strains, Rasika M. Harshey for the gift of the *Salmonella* strains, and Elizabeth Denson for creating the *Salmonella* typhimurium strain lacking the *tar* gene. We thank Dilara Long for contributing to simulations of an earlier version of the model.

## Supplementary Materials

**S1 Fig.**
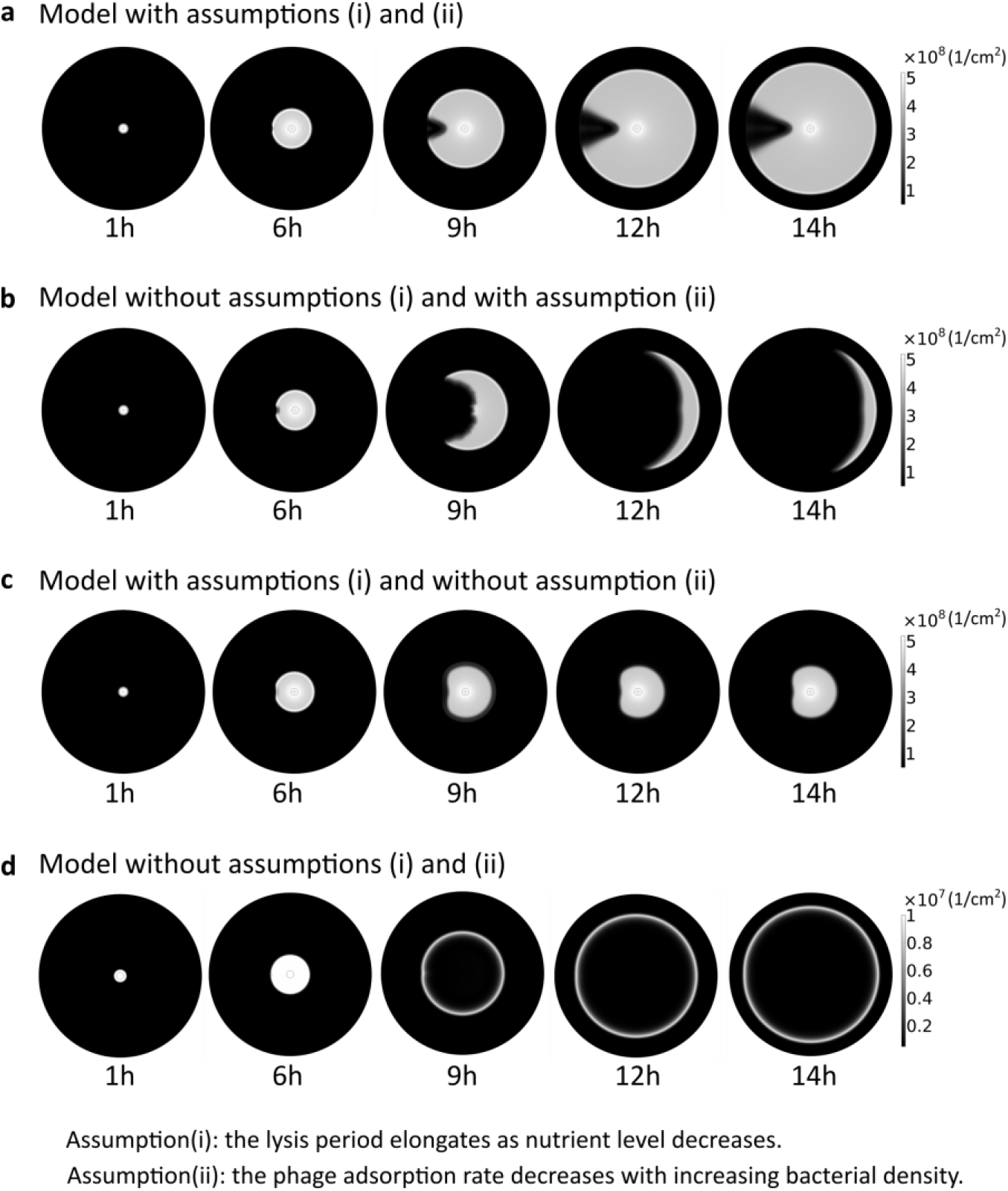
Both direct and indirect negative dependences of phage proliferation on bacterial density are necessary for generating straight radial boundaries in the lysis pattern. (a) Simulated lysis pattern formation with both Assumptions (i) and (ii). Same results as Fig. 1b, second row. (b) Simulated lysis pattern formation without Assumption (i), but with Assumption (ii). (c) Simulated lysis pattern formation without Assumption (ii), but with Assumption (i). (d) Simulated lysis pattern formation without both Assumptions. As described in Results, Assumption (i) states that nutrient deficiency inhibits phage replication, and Assumption (ii) states that high bacterial density inhibits phage production.

**S2 Fig.**
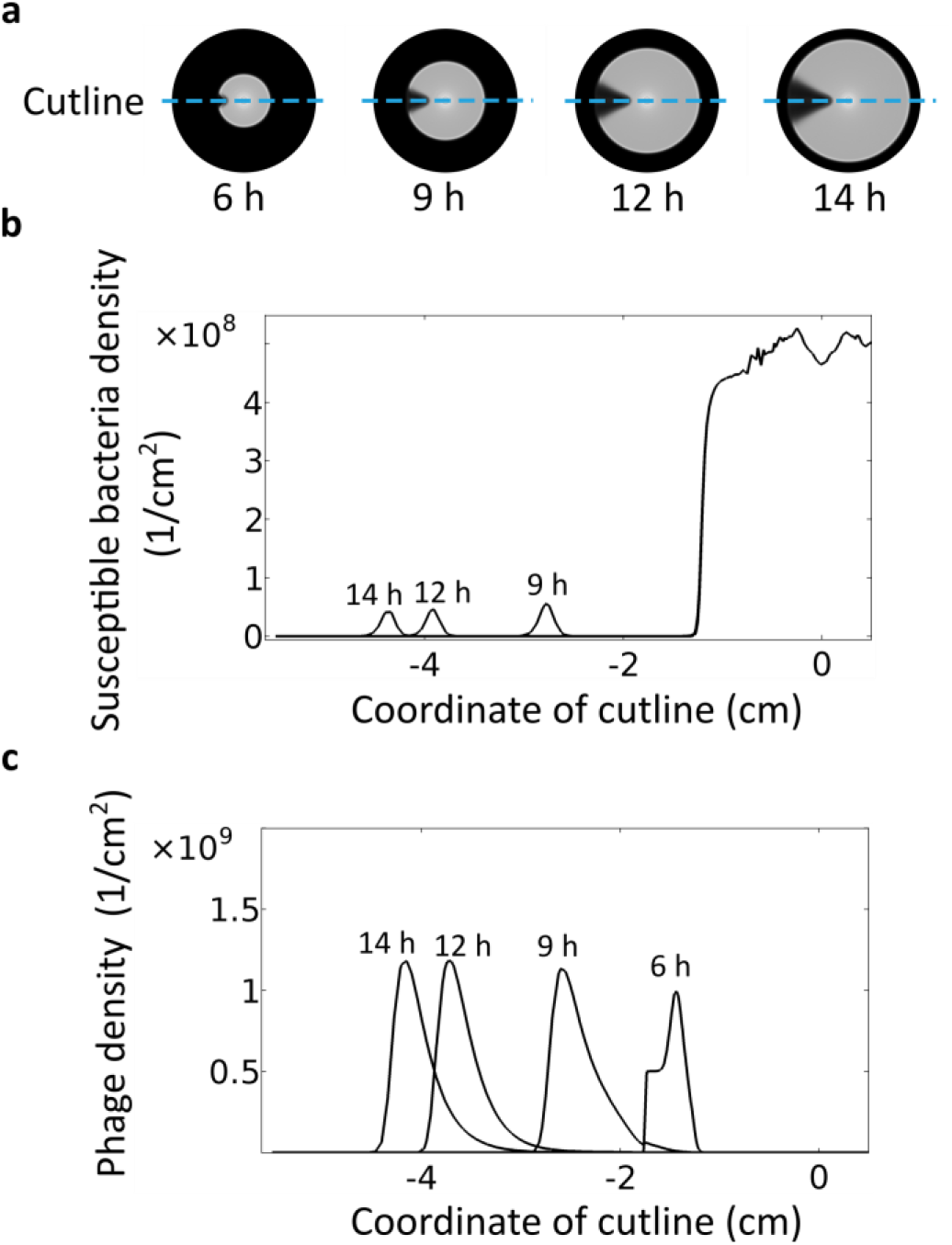
Model results show steady bacteria and phage densities at the expanding front after the phage initiation zone emerges. (a) Lysis patterns over time. Blue dashed line: cutline over which the density profiles are plotted in (b). (b) Density profiles of susceptible bacteria over the cutline at the labeled times. (c) Density profiles of phages over the cutline at the labeled times.

**S3 Fig.**
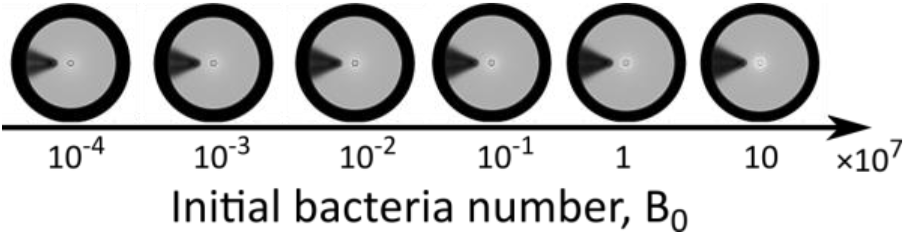
Simulation results with various numbers of inoculated bacteria.

**S4 Fig.**
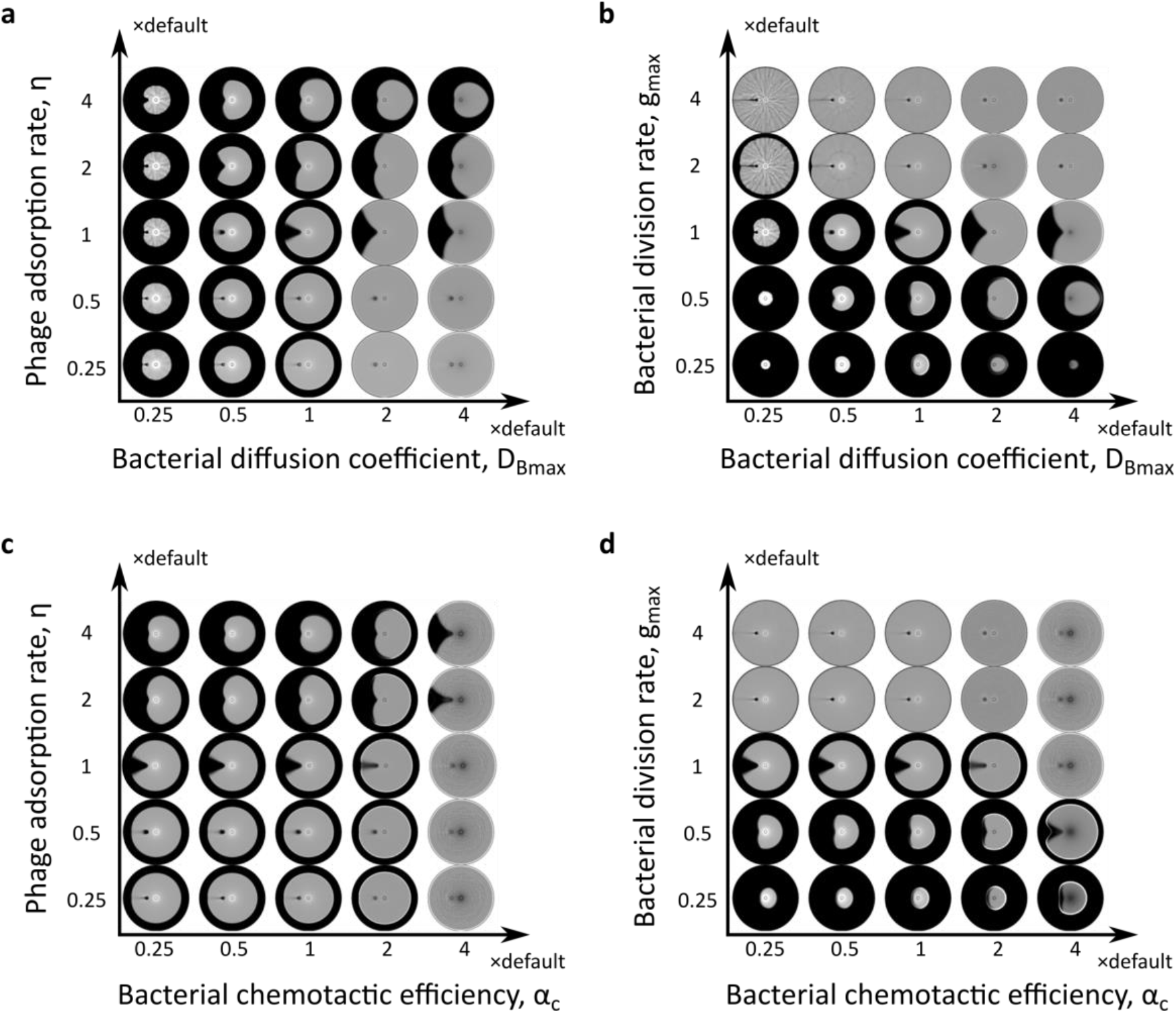
Lysis pattern with straight radial boundaries requires balance between intrinsic properties of bacteria and phage. Simulated lysis patterns with (a) various phage adsorption rate constants and bacterial diffusion coefficients, (b) various bacterial division rate constants and bacterial diffusion coefficients, (c) various phage adsorption rate constants and chemotactic efficiencies, and (d) various bacterial division rate constants and chemotactic efficiencies.

**S5 Fig.**
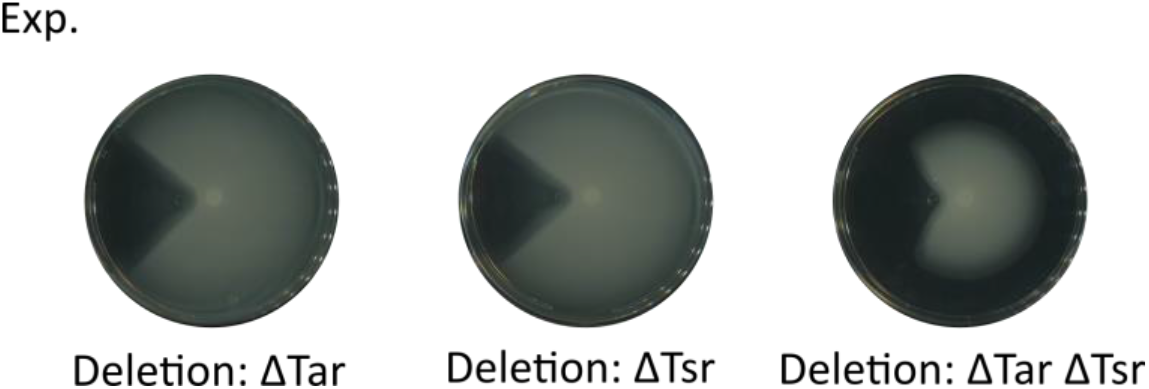
Experimental results of *S*. Typhimurium strains lacking chemoreceptors.

**S6 Fig.**
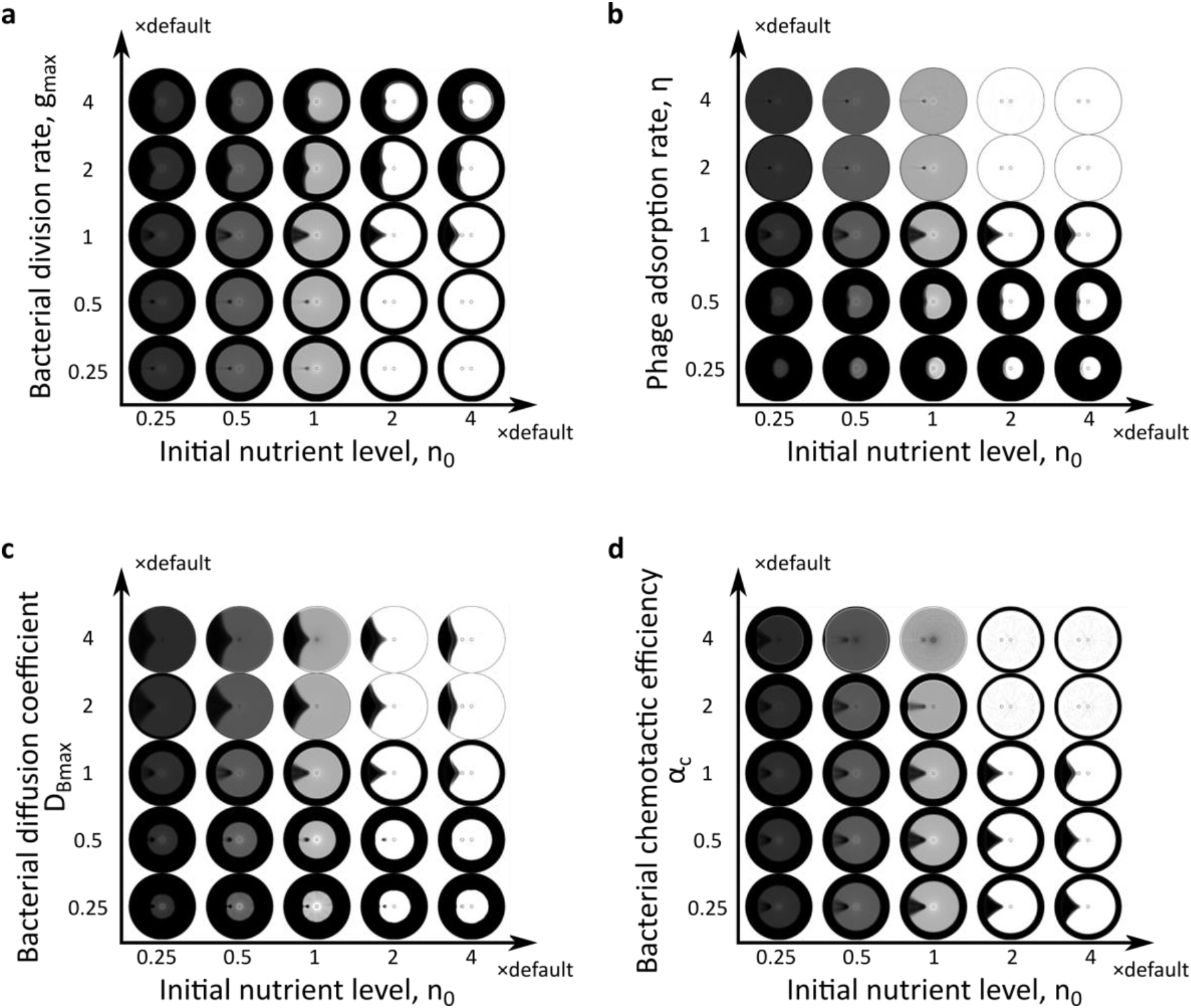
The shape of lysis area depends on the competition between phages and bacteria, but not the initial nutrient level. Simulated lysis patterns with various initial nutrient levels and (a) bacterial division rate constants, (b) phage adsorption rate constants, (c) bacterial diffusion coefficients, (d) chemotactic efficiencies.

**S1 Movie.**
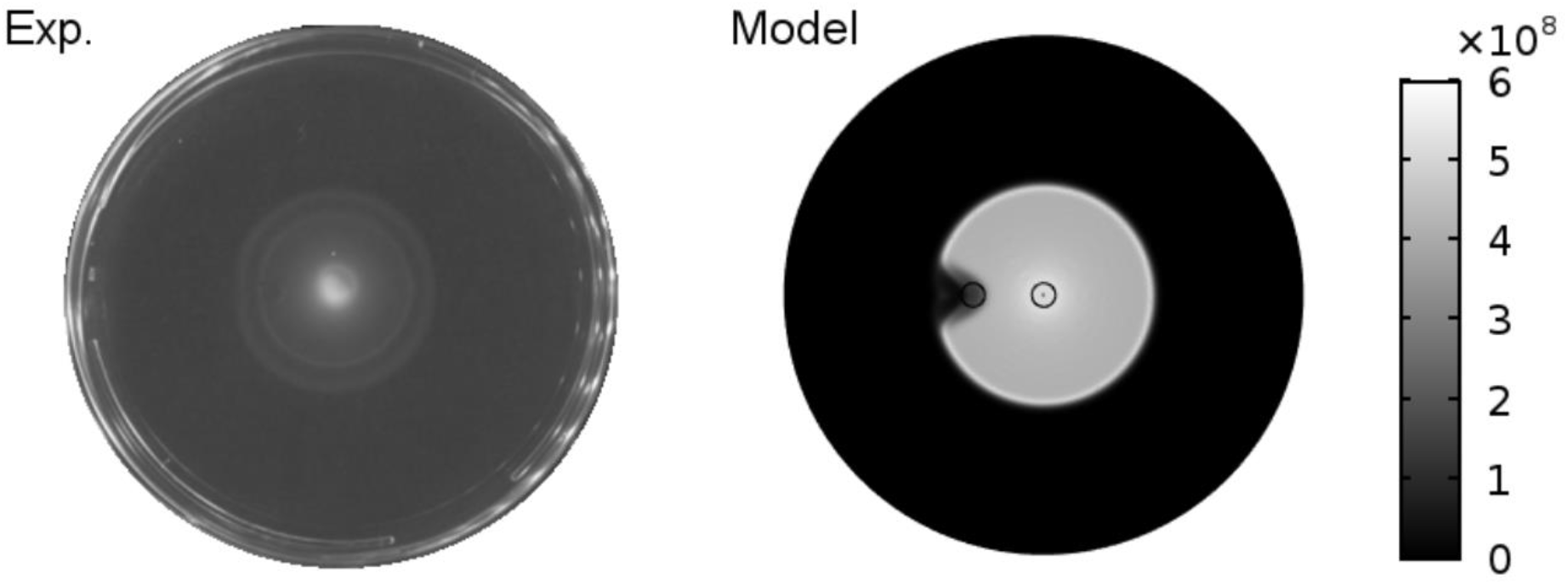
Time evolution of the lysis pattern in experiment and model. Each frame displays corresponding time points in experiment vs. model. Total time 14 hrs. Color bar represents density of bacteria (cm^−2^) in model.

### Mean field model for bacterial diffusion and chemotaxis with nutrient dependent motility

Here we derive the mean-field diffusion and chemotaxis terms, following the original work by Keller and Segel [1]. Specifically our model incorporates a nonlinear dependence of bacterial motility on nutrient level (as bacteria require energy to move). Note that both the diffusion and chemotaxis flux terms result from the run-and-tumble process of the bacteria; hence any regulation of bacterial motility or tumbling would affect both terms. Our derivation below emphasizes such quantitative relation between the diffusion and chemotaxis fluxes. For simplicity we derive the flux terms in 1D space, and the result can be readily generalized to 2D space. Our derivation is based on the following assumptions.

1. The bacterium takes left or right steps of length A.
2. The original paper by Keller and Segel assumes that the average frequency of steps to the left is governed by the mean nutrient concentration at the left edge, and vice versa [1]. Hence the difference in nutrient concentration across the cell body length governs the chemotaxis flux. This gradient sensing mode, however, differs from the temporal mode exploited by real bacteria [2]. Instead of detecting the difference of nutrient concentration at two ends of the cell (which is too small for reliable gradient detection in a tiny bacterial cell), the bacterial chemoreceptors use molecular memory to detect the temporal increase or decrease of nutrient signals as the bacterium swims [3]. However, note that the cell body length pictured in Keller and Segel’ original derivation essentially reflects the distance over which nutrient gradient is detected. Therefore, if we simply generalize the cell body to a “gradient-sensing distance”, the same derivation will apply to the temporal mode of gradient sensing (Fig. S7). In the temporal mode, the gradient-sensing distance reflects the distance a bacterium swims in the characteristic time of chemoreceptor memory.

**S7 Fig.**
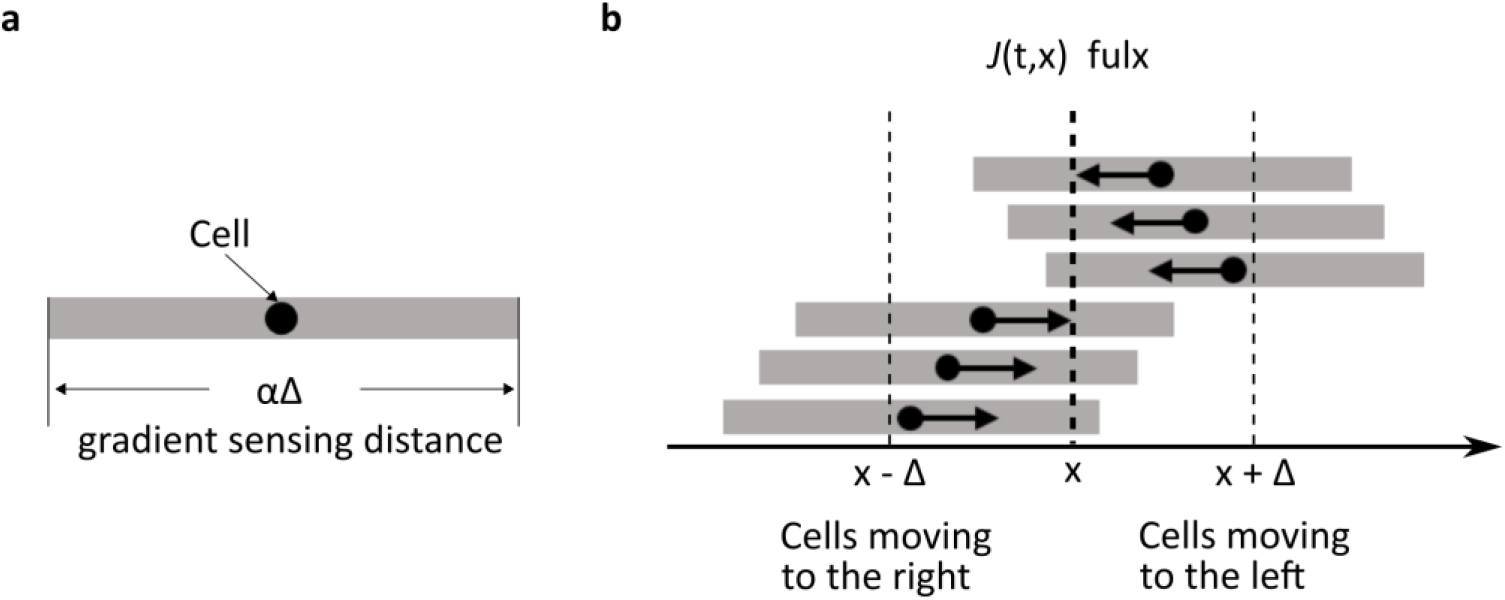
Spatial flux of bacteria. (a) Generalization of cell body length in the Keller-Segel derivation to gradient-sensing distance. (b) Illustration of the bacterial flux through a spatial location *x*. The flux of right-moving cells (lower batch) passing through the location *x* in a single step is the integration of the product of the step frequency to the right and cell density over the interval [*x* − Δ, *x*] (first integration term in Eq. (S1)). Vice versa, the flux of left-moving cells (upper batch) going through the location *x* in a single step is the integration of the product of the step frequency to the left and cell density over the interval [*x, x* + Δ] (second integration term in Eq. (S1)).

Let *B*(*t, x*) denote bacterial density at time t and location*x*. Let *f*(*n*) denote the average frequency of steps in a given direction, which is dependent on the local nutrient concentration, *n*(*t, x*). *α* is the ratio of the gradient-sensing distance to the step size. For a bacterium centered at location*x*, its frequencies of steps to the right and left, according to the 2nd assumption above, depend on the nutrient level at the right and left end of the gradient-sensing distance, i.e., written as 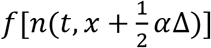 and 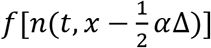, respectively. The net flux of bacteria *J*(*t, x*) at time *t* and location *x* (Fig. S9b) is defined by the amount of bacteria per unit time moving to the right minus the amount of bacteria per unit time moving to the left [1] (Eq.(S1)).

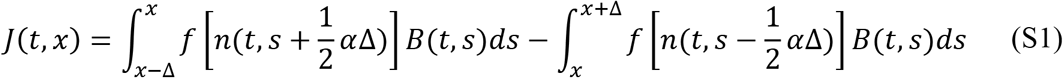

Note that when the step length Δ is sufficiently small, we can approximate the flux term up to *O*(*Δ*^2^), as shown in Eqs.(S2) and (S3).

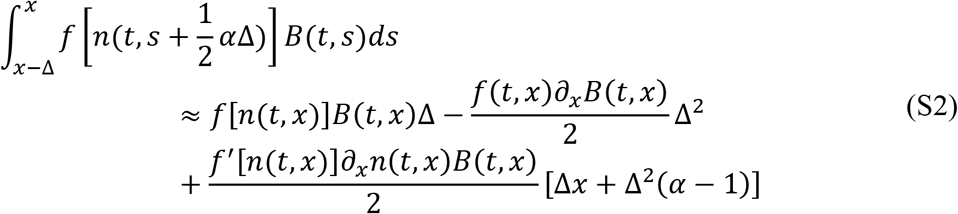

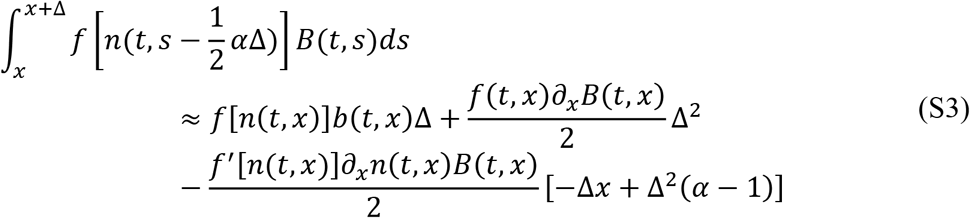

Plugging Eqs.(S2) and (S3) into Eq.(S1) yields

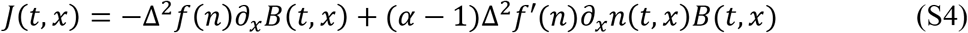

The first term describes diffusion (random motion) of the cells, and the second one describes the chemotaxis (biased motion up the nutrient gradient).

Because bacteria require energy to move, the cell velocity is likely an increasing function of nutrient level with a maximum value. We assumed a Michaelis-Menten type relation between nutrient level and cell velocity. Because *f*(*n*) is proportional to the cell velocity, it is also a Michaelis-Menten function of nutrient level (Eq. (S5)).

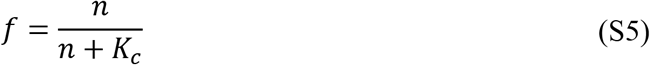

where*K_c_* is the nutrient level that allows half maximum cell velocity.

From Eq.(S4), we obtain the effective diffusion coefficient as a function of local nutrient level (Eq.(S6)).

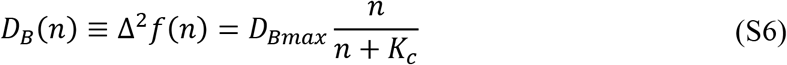

where the maximum diffusion coefficient, *D_Bmax_*, reflects maximum bacterial motility. From Eq.(S4) we also obtain the chemotactic coefficient *χ*(*n*) (Eq.(S7)).

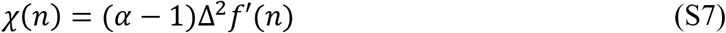

which can be rewritten as

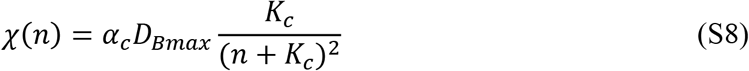

where*α_c_* = *α* − 1 represents the chemotactic efficiency. Note that the chemotactic coefficient given in Eq.(S8) is similar to that in the classic Keller-Segel models [4]. However, instead of representing a saturated chemotactic response, the fraction term in Eq.(S8) represents the saturation of cell velocity, with increasing nutrient level, as a related fraction term also appears in the diffusion flux (Eq.(S6)).

The overall spatial flux of bacteria in 2D space, including random motion and chemotaxis process, is derived by replacing the 1D spatial derivative, *∂_x_*, by the 2D gradient operator, ∇=[*∂_x_, ∂_y_*] (Eq.S(9)).

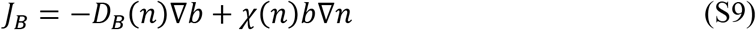

